# History of childbirths relates to region-specific brain aging patterns in middle and older-aged women

**DOI:** 10.1101/2020.05.08.084616

**Authors:** Ann-Marie G. de Lange, Claudia Barth, Tobias Kaufmann, Melis Anatürk, Sana Suri, Klaus P. Ebmeier, Lars T. Westlye

**Affiliations:** Department of Psychiatry, University of Oxford, Oxford, UK; Department of Psychology, University of Oslo, Oslo, Norway; NORMENT, Institute of Clinical Medicine, University of Oslo, & Division of Mental Health and Addiction, Oslo University Hospital, Oslo, Norway; Wellcome Centre for Integrative Neuroimaging, University of Oxford, Oxford, UK; KG Jebsen Centre for Neurodevelopmental Disorders, University of Oslo, Oslo, Norway

## Abstract

Pregnancy involves maternal brain adaptations, but little is known about how parity influences women’s brain aging trajectories later in life. In this study, we replicated previous findings showing less apparent brain aging in women with a history of childbirths, and identified regional brain aging patterns linked to parity in 19,787 middle and older-aged women. Using novel applications of brain-age prediction methods, we found that a higher number of previous childbirths was linked to less apparent brain aging in striatal and limbic regions. The strongest effect was found in the accumbens – a key region in the mesolimbic reward system, which plays an important role in maternal behavior. While only prospective longitudinal studies would be conclusive, our findings indicate that subcortical brain modulations during pregnancy and postpartum may be traceable decades after childbirth.

## Introduction

Pregnancy involves a number of maternal brain adaptations (***Barha and Galea, 2017***; ***Fox et al., 2018***; ***Hillerer et al., 2014***; ***Eid et al., 2019***; ***Boddy et al., 2015***). In rodents, changes in volume, cell proliferation, and dendritic morphology (***Hillerer et al., 2014***; ***Kinsley et al., 2006***), as well as altered neurogenesis in the hippocampus (***Eid et al., 2019***; ***Rolls et al., 2008***) are found across pregnancy and postpartum. In humans, reduction in total brain volume has been observed during pregnancy, reversing within six months of parturition (***Oatridge et al., 2002***). Reductions in striatal volumes, particularly putamen, have been reported shortly after delivery (***Lisofsky et al., 2016***), and pregnancy-related reductions in gray matter volume have been found in regions subserving social cognition; the bilateral lateral prefrontal cortex, the anterior and posterior midline, and the bilateral temporal cortex (***Hoekzema et al., 2017***). Conversely, a recent study showed no evidence of decrease in gray matter volume following childbirth, but instead detected a pronounced gray matter *increase* in both cortical and subcortical regions (***Luders et al., 2020***). Prefrontal cortical thickness and subcortical volumes in limbic areas have been positively associated with postpartum months (***Kim et al., 2018***), indicating that changes in brain structure may depend on region and time since delivery (***Luders et al., 2020***; ***Duarte-Guterman et al., 2019***; ***Hoekzema et al., 2017***; ***Kim et al., 2018, 2010***). For instance, from 2–4 weeks to 3–4 months postpartum, gray matter volume increases have been found in areas involved in maternal behaviours and motivation, such as the amygdala, substantia nigra, hypothalamus, and prefrontal cortex (***Kim et al., 2010***).

While gray matter reductions have been reported up to 2 years post-pregnancy, most studies are limited to the postpartum period, and little is known about how previous pregnancies influence women’s brain aging later in life. Evidence from animal studies shows that middle-aged multiparous rats have stronger cellular response to estrogens in the hippocampus compared to virgin female rats (***Barha and Galea, 2011***), suggesting that neuroplastic potential across the adult lifespan may be influenced by previous pregnancies. Moreover, hippocampal neurogenesis has been shown to increase during middle age in primiparous rats and decrease in nulliparous rats over the same period (***Eid et al., 2019***). While longitudinal studies on parity and brain aging in humans are lacking, cumulative number of months pregnant has been associated with decreased risk for Alzheimer’s disease (***Fox et al., 2018***), and we recently documented less evident brain aging in parous relative to nulliparous women in >12,000 UK Biobank participants using an magnetic resonance imaging (MRI)-derived biomarker of global brain aging (***de Lange et al., 2019***).

In the current study, we first aimed to replicate our previously reported findings described in ***de Lange et al***. (***2019***), where less apparent brain aging was found in women with a history of childbirths. Brain-age prediction methods were used to derive estimates of global brain aging, which was analysed in relation to number of previous (live) childbirths in 8880 newly added UK Biobank participants. Brain-age prediction is commonly used to estimate an individual’s age based on their brain characteristics (***Cole and Franke, 2017***), and individual variation in “brain age” estimates has been associated with a range of clinical and biological factors (***Cole and Franke, 2017***; ***Kaufmann et al., 2019***; ***Franke and Gaser, 2019***; ***Cole et al., 2018, 2019***; ***Cole, 2019***; ***Smith et al., 2019***; ***Cole et al., 2017***; ***de Lange et al., 2020a***,b). As compared to MRI-derived measures such as cortical volume or thickness, brain-age prediction adds a dimension by capturing deviations from normative trajectories identified by machine learning. While traditional brain age approaches summarize measures across regions to produce a single, global aging estimate – often with high prediction accuracy, models of distinct and regional aging patterns can provide more refined biomarkers that may capture additional biological detail (***Kaufmann et al., 2019***; ***Smith et al., 2020***; ***Eavani et al., 2018***). In this study, we utilized novel applications of brain-age prediction methods based on cortical and subcortical volumes to identify regions of particular importance for maternal brain aging, using a total sample of 19,787 UK Biobank women.

## Results

### Associations between previous childbirths and global brain aging

To replicate our findings described in ***de Lange et al***. (***2019***), we trained a brain-age prediction model on the part of the current sample that overlapped with the previous study (N = 10,907), and applied it to the newly added participants (N = 8880) using the procedure described in *Methods and Materials*. When applied to the test set, the modeled age prediction showed an accuracy of R^2^ = 0.34, root mean square error (RMSE) = 6.00, and Pearson’s r (predicted versus chronological age) = 0.58; 95% confidence interval (CI) = [0.57, 0.59], *p* < 0.001. Corresponding to our previous results, an association was found between a higher number of previous childbirths and less apparent brain aging in the group of newly added participants: *β* = −0.13 standard error (SE) = 0.03, *t* = −4.07, *p* = 4.79×10^−5^. To adjust for age-bias in the brain age prediction as well as age-dependencies in number of childbirths (***Le et al., 2018***), chronological age was included as a covariate in the linear regression.

To test for non-linear relationships, polynomial fits were run for number of childbirths and brain age gap; one including intercept and a linear term (*β*) only, and one including intercept, linear, and quadratic terms (*γ*). For these analyses, the brain age gap estimations were first corrected for chronological age using linear regression (***Le et al., 2018***), and the residuals were used in the fits. A comparison of the two models showed that the inclusion of the quadratic term did not provide a better fit (*F* = 0.06, *p* = 0.804). The results from the fit including the linear term only showed a significant linear effect (*β* = −0.12 ± 0.03; *F* = 16.10, *p* = 6.05 × 10^−5^), while the results from the fit including both terms showed that only the linear term was significant (*β* = −0.14 ± 0.072; *γ* = 0.004 ± 0.02; *F* = 8.08, *p* = 3.11 × 10^−4^). The two fits are shown in Figure 1. As a cross check, the fits were rerun with orthogonal polynomials, showing corresponding results (*β* = −13.44 ± 3.35, *γ* = 0.83± 3.35; *F* = 8.08, *p* = 3.11× 10^−4^).

**Figure 1.**
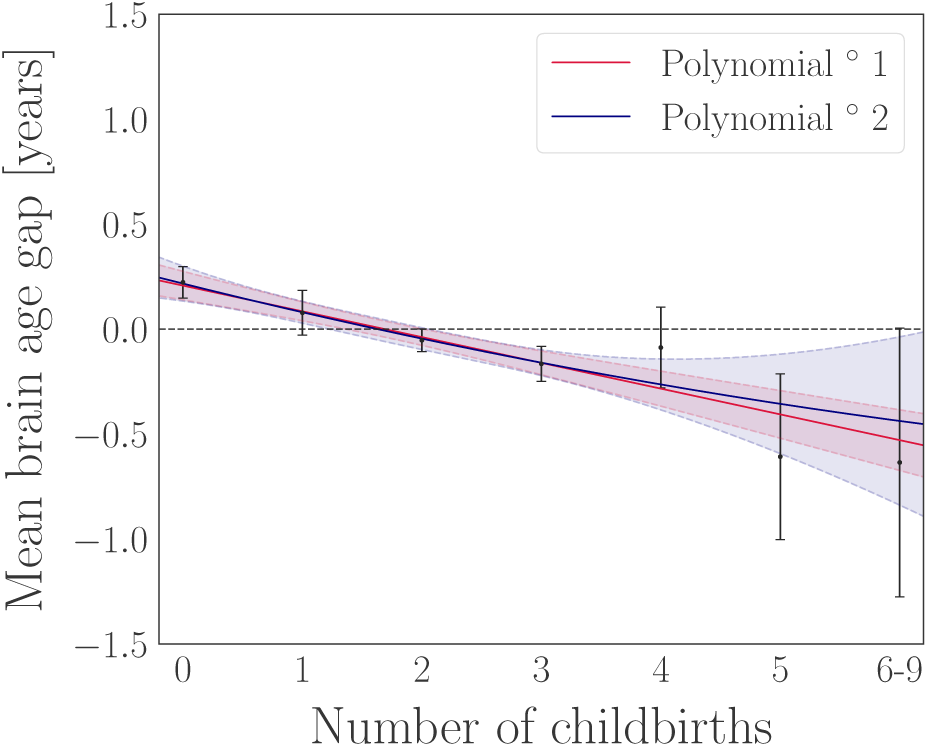
Results from first and second degree polynomial fits for number of childbirths and global brain aging in the newly added participants (N = 8880). The black points indicate the mean brain age gap ± standard error within groups of women based on number of childbirths (x-axis). The red and blue lines represent the results of the fits, and the shaded areas indicate the ± 95% confidence intervals for each fit. The horizontal dashed line indicates 0 on the y-axis. Number of participants in each group: 0 births = 2065, 1 birth = 1014, 2 births = 3912, 3 births = 1493, 4 births = 311, 5 births = 67, 6 births = 13, 7 births = 3, 8 births = 1, and 9 births = 1. The group of women with 6-9 children were merged to obtain sufficient statistics for least square fits using the standard error on the means as weights.

### Regional brain aging patters

To identify groups of imaging features based on their covariance, we performed hierarchical clustering on the Spearman rank-order correlations using an average of right and left hemisphere measures for each feature. Five clusters were identified, as shown in Figure 2. All features were grouped according to their associated cluster ID, and separate models were run to estimate brain age for each cluster using the brain-age prediction procedure described in *Methods and Materials*. The features contained in each cluster are listed in Table 1. The cluster-specific model performances are shown in Table 2.

**Table 1.** List of imaging features contained in each of the clusters identified based on hierarchical clustering (Figure 2). All feature names represent regional volume.

|  |  |  |  |
| --- | --- | --- | --- |
| **Cluster 1** |  |  |  |
| Cuneus | Isthmuscingulate | Lateraloccipital | Lingual |
| Pericalcarine | Precuneus | Superiorparietal |  |
| **Cluster 2** |  |  |  |
| Caudalanteriorcingulate | Lateralorbitofrontal | Medialorbitofrontal | Paracentral |
| Parsopercularis | Parsorbitalis | Parstriangularis | Postcentral |
| Posteriorcingulate | Precentral | Rostralmiddlefrontal | Superiorfrontal |
| Superiortemporal | Supramarginal | Transversetemporal | Insula |
| **Cluster 3** |  |  |  |
| Banks of superior temporal sulcus | Fusiform | Inferiorparietal | Inferiortemporal |
| Middletemporal | Parahippocampal | Thalamus | Putamen |
| Hippocampus | Amygdala | Accumbens |  |
| **Cluster 4** |  |  |  |
| Caudalanteriorcingulate | Entorhinal | Rostralanteriorcingulate | Frontalpole |
| Temporalpole |  |  |  |
| **Cluster 5** |  |  |  |
| Brain-Stem | Cerebellum | Caudate | Pallidum |

**Table 2.** The accuracy of the age prediction measured by Pearson’s r (predicted versus chronological age), R^2^, root mean square error (RMSE), and mean absolute error (MAE) for each of the cluster-specific models. 95 % confidence intervals are indicated in square brackets. RMSE and MAE are reported in years. *N*_*feat*_ = number of features contained in the cluster. *P*-values were < 0.001 for all models.

| Model | $N_{feat}$ | $r$ | $R^2$ | RMSE | MAE |
| --- | --- | --- | --- | --- | --- |
| Cluster 1 | 7 | 0.32 [0.31, 0.33] | 0.10 [0.10, 0.11] | 7.00 | 5.79 |
| Cluster 2 | 16 | 0.38 [0.36, 0.39] | 0.14 [0.13, 0.15] | 6.86 | 5.64 |
| Cluster 3 | 11 | 0.48 [0.47, 0.49] | 0.23 [0.23, 0.24] | 6.48 | 5.29 |
| Cluster 4 | 5 | 0.13 [0.11, 0.14] | 0.02 [0.01, 0.02] | 7.36 | 6.13 |
| Cluster 5 | 4 | 0.21 [0.19, 0.22] | 0.04 [0.04, 0.05] | 7.25 | 6.04 |

**Figure 2.**
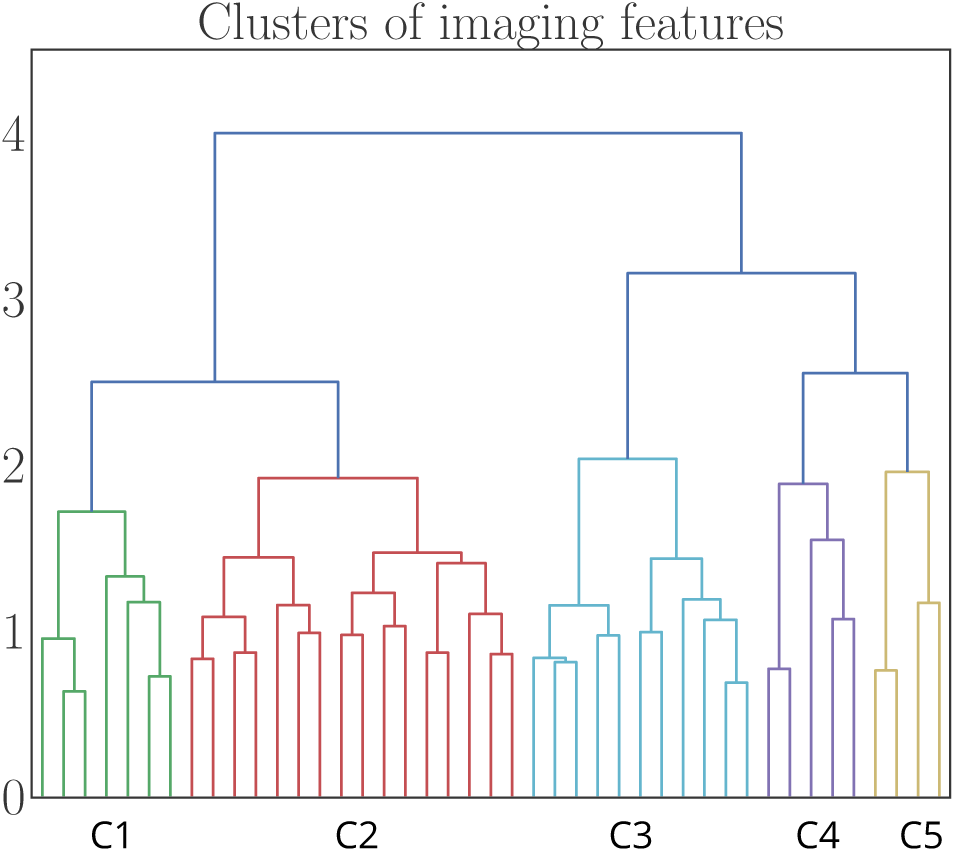
Dendrogram based on hierarchical clustering on the Spearman rank-order correlations of all features. The colours represent clusters (C) of features that are grouped together based on common covariance. A list of imaging features contained in each of the clusters are provided in Table 1. The y-axis shows the degree of co-linearity between the features, with higher y-values indicating less co-linearity between clusters.

To test whether the relative prediction accuracy of the models depended on number of features, the models were rerun using the four strongest contributing features from each model as input variables. The feature contributions were calculated using permutation feature importance, defining the decrease in model performance when a single feature value is randomly shuffled (***Breiman, 2001***). The results are shown in Table 3.

**Table 3.** The accuracy of the age prediction when including the four strongest contributing features for each model. 95 % confidence intervals are indicated in square brackets. RMSE and MAE are reported in years. *N*_*feat*_ = number of features included. *P*-values were < 0.001 for all models.

| Model | $N_{feat}$ | $r$ | $R^2$ | RMSE | MAE |
| --- | --- | --- | --- | --- | --- |
| Cluster 1 | 4 | 0.32 [0.31, 0.33] | 0.10 [0.10, 0.11] | 7.01 | 5.80 |
| Cluster 2 | 4 | 0.34 [0.33, 0.35] | 0.11 [0.11, 0.12] | 6.96 | 5.75 |
| Cluster 3 | 4 | 0.46 [0.45, 0.47] | 0.21 [0.21, 0.22] | 6.55 | 5.36 |
| Cluster 4 | 4 | 0.12 [0.11, 0.13] | 0.02 [0.01, 0.02] | 7.36 | 6.14 |
| Cluster 5 | 4 | 0.21 [0.19, 0.22] | 0.04 [0.04, 0.05] | 7.25 | 6.04 |

### Associations between previous childbirths and regional brain aging

The cluster-specific associations with number of previous childbirths are shown in Table 4.

**Table 4.** Relationships between number of previous childbirths and estimated brain aging for each cluster. Cluster-specific *brain age gap* was entered as the dependent variable and *number of (live) childbirths* was entered as independent variable for each analysis. Chronological age was included for covariate purposes. *P*-values are provided before and after FDR correction. Significant relationships (< 0.05) are marked with an asterisk in the *p*_*corr*_ column. SE = standard error.

| Number of childbirths vs cluster-specific brain age estimates |  |  |  |  |  |
| --- | --- | --- | --- | --- | --- |
| Cluster | $\beta$ | SE | $t$ | $p$ | $p_{corr}$ |
| 1 | -0.054 | 0.016 | -3.282 | 0.001 | 0.002* |
| 2 | -0.010 | 0.018 | -0.552 | 0.581 | 0.726 |
| 3 | -0.133 | 0.021 | -6.385 | $1.754 \times 10^{-10}$ | $8.768 \times 10^{-10}$* |
| 4 | -0.003 | 0.010 | -0.278 | 0.781 | 0.781 |
| 5 | -0.056 | 0.012 | -4.620 | $3.868 \times 10^{-16}$ | $9.670 \times 10^{-6}$* |

As shown in Table 4, brain aging estimates based on three clusters were each significantly associated with number of previous childbirths. To test whether the associations were statistically different from each other, pairwise Z tests for correlated samples (Eq. 1, *Methods and Materials*) were run on the cluster-specific associations with number of childbirths. The results showed that cluster 3 was more strongly related to number of previous childbirths relative to the other clusters, as shown in Figure 3.

**Figure 3.**
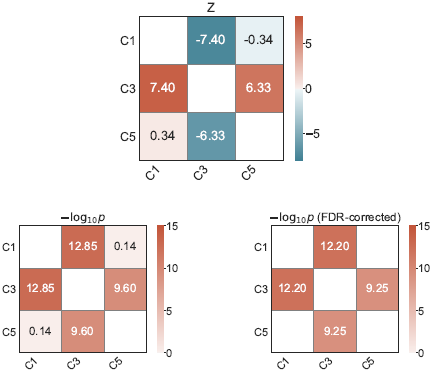
Top plot: Matrix showing pairwise differences between the cluster-specific associations with number of childbirths, based on Z tests for correlated samples (Eq.1). Bottom left plot: Uncorrected −Log10 p-values of the differences between the cluster-specific associations. Bottom right plot: −Log10 *p*-values corrected for multiple comparisons using FDR, with only significant values (< 0.05) displayed. C = cluster.

To investigate further specificity, the clustering procedure was repeated on the features in cluster 3 – the cluster showing the strongest association with number of childbirths. Two subclusters were identified based on the covariance of the features, as shown in Figure 4. Subcluster 1 included 5 features; subcluster 2 included 6 features. The features were matched with the cluster they belonged to, and separate models were run to generate brain-age predictions for each subcluster. The subcluster-specific model performances are shown in Table 5, and their associations with number of previous childbirths are shown in Table 6.

**Table 5.** The accuracy of the age prediction measured by Pearson’s r (predicted versus chronological age), R^2^, root mean square error (RMSE), and mean absolute error (MAE) for each of the subcluster-specific models. 95 % confidence intervals are indicated in square brackets. RMSE and MAE are reported in years. *N*_*feat*_ = number of features contained in the cluster. P-values were < 0.001 for both models.

| Model | $N_{feat}$ | $r$ | $R^2$ | RMSE | MAE |
| --- | --- | --- | --- | --- | --- |
| Subcluster 1 | 5 | 0.33 [0.31, 0.34] | 0.11 [0.10, 0.12] | 6.99 | 5.79 |
| Subcluster 2 | 6 | 0.47 [0.45, 0.48] | 0.22 [0.20, 0.23] | 6.54 | 5.35 |

**Table 6.** Relationships between number of previous childbirths and estimated brain aging for each subcluster. Subcluster-specific *brain age gap* was entered as the dependent variable and *number of childbirths* was entered as independent variable for each analysis. Chronological age was included for covariate purposes. *P*-values are provided before and after FDR correction.

| Number of childbirths vs subcluster-specific brain age estimates |  |  |  |  |  |
| --- | --- | --- | --- | --- | --- |
| Subcluster | $\beta$ | SE | *t* | *p* | *P<sub>corr</sub>* |
| 1 | -0.071 | 0.016 | -4.321 | $1.563 \times 10^{-5}$ | $1.563 \times 10^{-5}$ |
| 2 | -0.124 | 0.020 | -6.169 | $7.024 \times 10^{-10}$ | $1.405 \times 10^{-9}$ |

**Figure 4.**
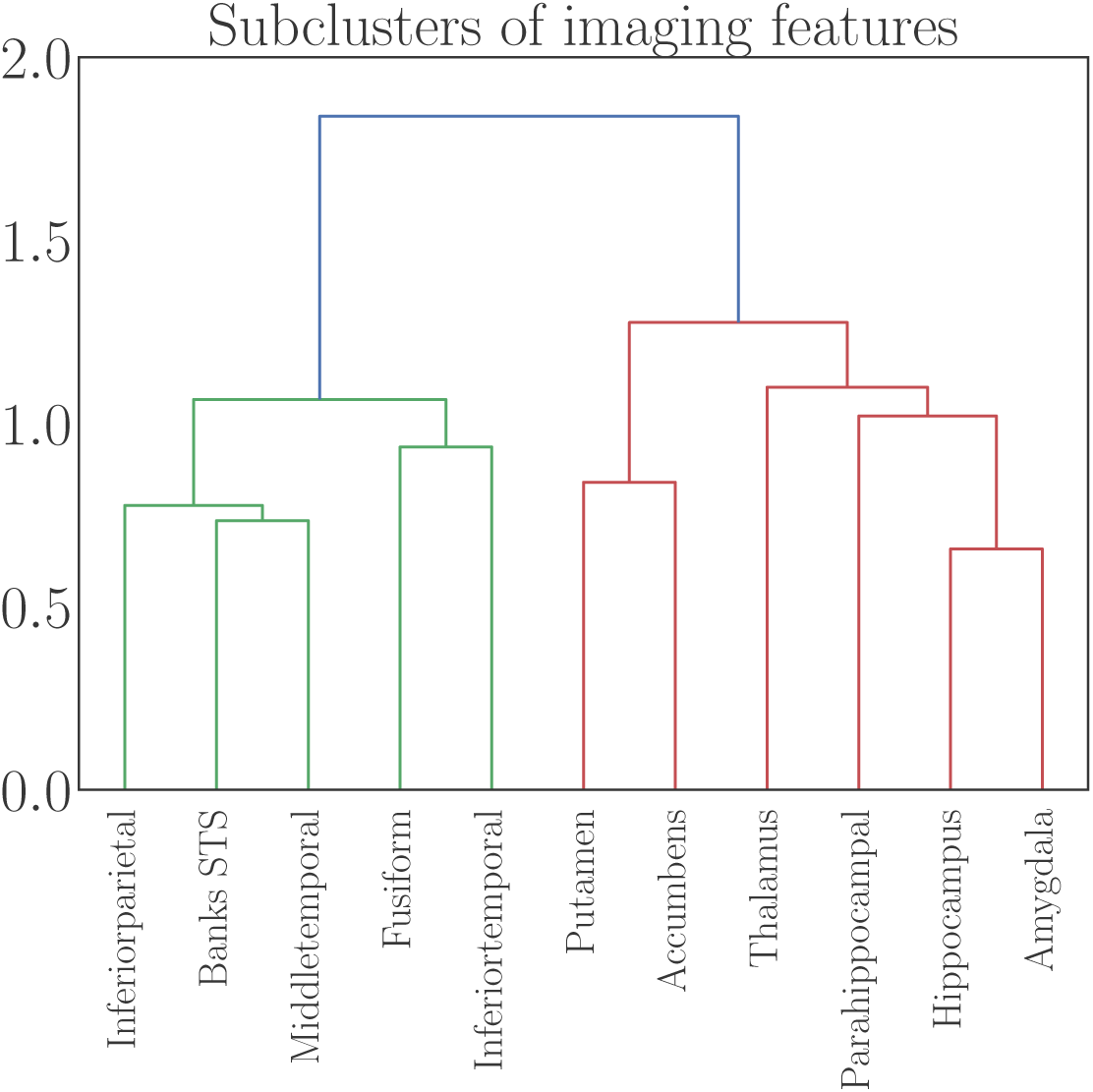
Dendrogram based on hierarchical clustering on the Spearman rank-order correlations of the features contained in Cluster 3, which showed the strongest association with number of childbirths (see Figure 3). The colours represent clusters of features that are grouped together based on common covariance; subcluster 1 in green and subcluster 2 in red. The y-axis shows the degree of co-linearity between the features, with higher y-values indicating less co-linearity between clusters. STS = superior temporal sulcus.

**Figure 5.**
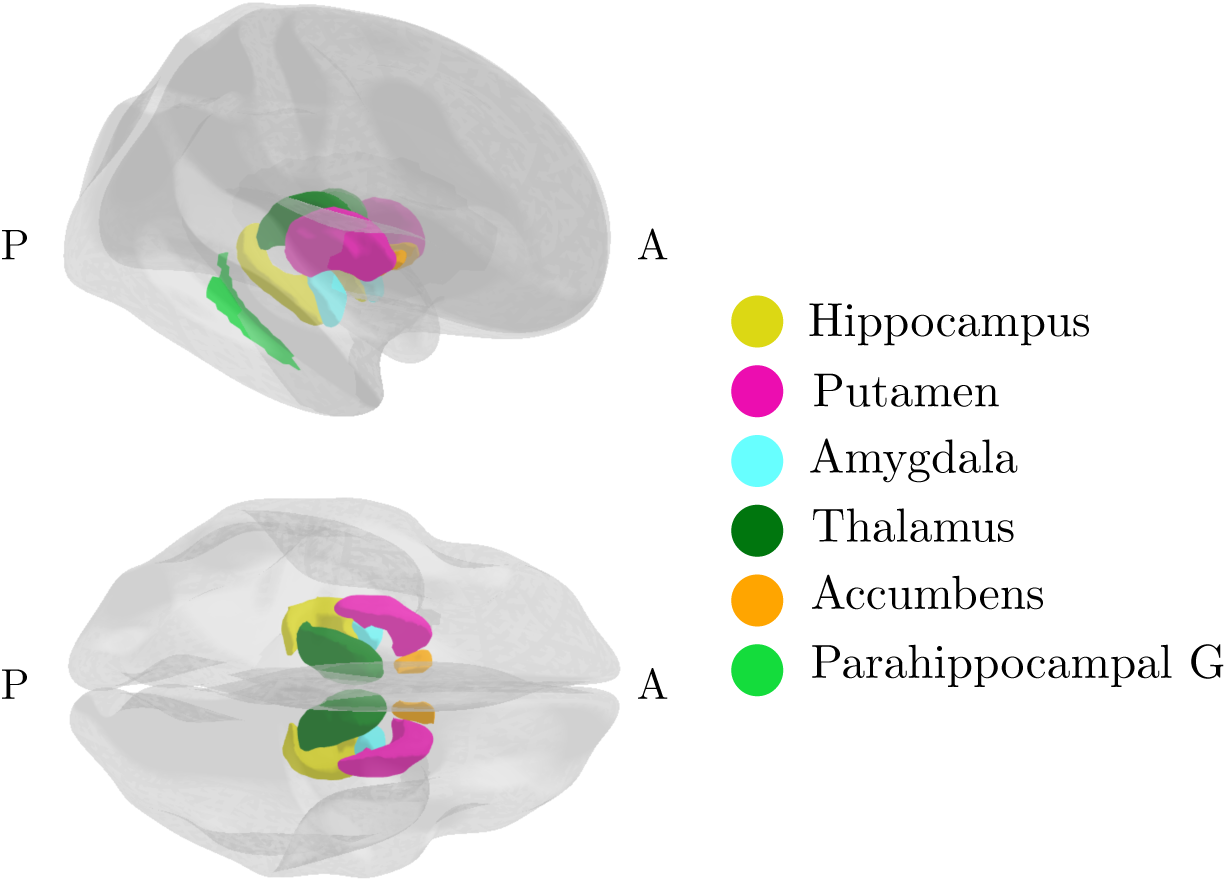
Regions in subcluster 2 – the cluster that showed the strongest association with number of previous childbirths. A = anterior, G = gyrus, P = posterior. Figure created using the *ggseg* plotting tool for brain atlases inR (***Mowinckel and Vidal-Piñeiro, 2019***).

To control for potential confounding factors, the analyses of number of previous childbirths versus subcluster 2 brain age estimates were rerun including assessment location, education, body mass index (BMI), diabetic status, hypertension, smoking and alcohol intake, menopausal status (‘yes’, ‘no’, ‘not sure, had hysterectomy’, and ‘not sure, other reason’), and oral contraceptive (OC) and hormonal replacement therapy (HRT) status (previous or current user vs never used) as covariates. 16,512 women had data on all variables and were included in the analyses. The results showed an association of *β* = −0.12, SE = 0.02, *t* = −5.36, *p* = 8.27 × 10^−8^ between number of childbirths and subcluster 2, indicating that the covariates could not fully explain the association. Number of previous childbirths and age at first birth correlated *r* = −0.294, *p* = 6.90× 10^−296^ (corrected for age). To test for an association with brain aging, an analysis was run with subcluster 2 brain age as the dependent variable and *age at first birth* as independent variable, including all the covariates (age, assessment location, education, BMI, diabetic status, hypertension, smoking and alcohol intake, menopausal status, OC and HRT use). No association was found (*β* = 0.010, SE = 0.01, *t* = 1.64, *p* = 0.102, N = 12, 937).

### Single-region associations

To investigate the unique contributions of each region in subcluster 2 to the association with previous childbirths, separate brain-age prediction models were run with each feature as input, yielding 11 region-specific brain age estimates. Table 8 shows the correlation between predicted and chronological age for each region-specific model, and their associations with number of childbirths. As the regions within the cluster were correlated (see Figure 4), we tested for unique contributions by first running a multiple regression analysis with all region-specific brain age estimates as independent variables and *number of childbirths* as the dependent variable, before eliminating the regions one at a time to compare the log-likelihood of the full and reduced models. The significance of model differences was calculated using Wilk’s theorem (***Wilks, 1938***) as 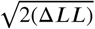, where Δ*LL* = *LL*_1_ − *LL*_2_; the difference in log-likelihood between the reduced model (*LL*_1_) and the full model (*LL*_2_). The results showed that only the accumbens contributed uniquely to the association with number of previous childbirths. The association when excluding accumbens from subcluster 2 was *β* = −0.098, SE = 0.020, *t* = −5.001, *p* = 5.766 × 10^−07^, indicating that the association was not solely driven by this region.

**Table 7.** Difference in quadrature between the subcluster-specific associations with number of childbirths, calculated using Eq. 1 (*Methods and Materials*).

| Comparison | *Z* | *p* | *P<sub>corr</sub>* |
| --- | --- | --- | --- |
| Subcluster 1 vs subcluster 2 | -5.32 | $1.01 \times 10^{-7}$ | $2.03 \times 10^{-7}$ |

**Table 8.** Region-specific age prediction accuracy (correlation between predicted and chronological age; *r*_*Age*_) and association with number of childbirths (*β*_*CB*_, standard error (SE), *t, p*, and *p*_*corr*_) for each of the region-specific brain age gap estimates. Chronological age was included in the analyses for covariate purposes. 95 % confidence intervals are indicated in square brackets. *P*-values are reported before and after FDR correction.

| Subcluster 2 region | $r_{Age}$ | $\beta_{CB}$ | SE | $t$ | $p$ | $p_{corr}$ |
| --- | --- | --- | --- | --- | --- | --- |
| Parahippocampal | 0.24 [0.23, 0.25] | -0.020 | 0.012 | -1.650 | 0.099 | 0.099 |
| Thalamus | 0.35 [0.34, 0.36] | -0.048 | 0.016 | -2.991 | 0.003 | 0.003 |
| Putamen | 0.24 [0.22, 0.25] | -0.053 | 0.012 | -4.401 | $1.08 \times 10^{-5}$ | $2.253 \times 10^{-5}$ |
| Hippocampus | 0.33 [0.32, 0.34] | -0.061 | 0.016 | -3.923 | $8.77 \times 10^{-5}$ | $1.315 \times 10^{-4}$ |
| Amygdala | 0.29 [0.27, 0.30] | -0.061 | 0.014 | -4.393 | $1.13 \times 10^{-5}$ | $2.252 \times 10^{-5}$ |
| Accumbens | 0.31 [0.30, 0.32] | -0.101 | 0.015 | -6.812 | $9.90 \times 10^{-12}$ | $5.937 \times 10^{-11}$ |

As a cross check, we investigated associations between previous childbirths and regional volumes in subcluster 2. Separate analyses were run with the volume measure for each region as dependent variables and number of previous childbirths as the independent variable, including age, assessment location, education, BMI, diabetic status, hypertension, smoking and alcohol intake, menopausal status, and OC and HRT status as covariates. 16,516 women had data on all variables and were included in the analysis. The associations between number of previous childbirths and regional volume corresponded to the associations with brain age estimates, as shown in Table 10.

**Table 9.**
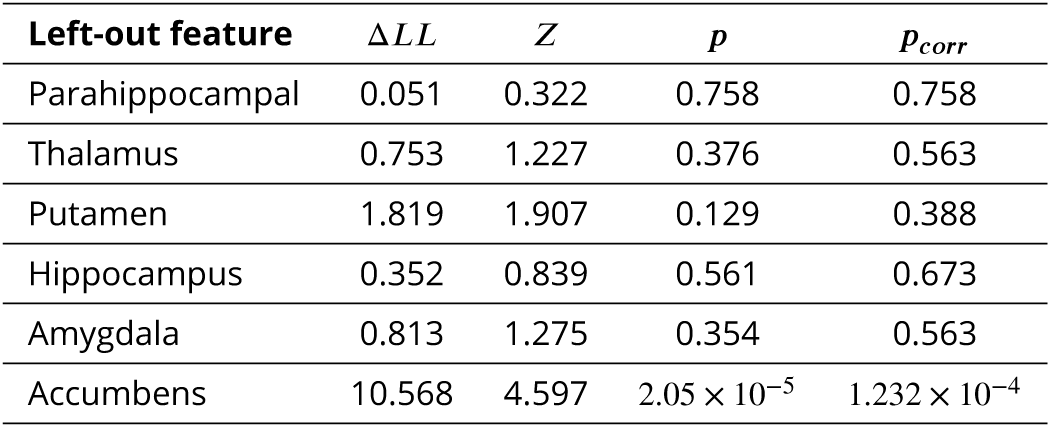
Difference in log-likelihood (Δ*LL*) between regression analyses against *number of children*. The difference is calculated between models where all cluster features are included and models where single features are left out one at the time. *P*-values are reported before and after FDR correction.

| Left-out feature | $\Delta LL$ | $Z$ | $p$ | $p_{corr}$ |
| --- | --- | --- | --- | --- |
| Parahippocampal | 0.051 | 0.322 | 0.758 | 0.758 |
| Thalamus | 0.753 | 1.227 | 0.376 | 0.563 |
| Putamen | 1.819 | 1.907 | 0.129 | 0.388 |
| Hippocampus | 0.352 | 0.839 | 0.561 | 0.673 |
| Amygdala | 0.813 | 1.275 | 0.354 | 0.563 |
| Accumbens | 10.568 | 4.597 | $2.05 \times 10^{-5}$ | $1.232 \times 10^{-4}$ |

**Table 10.**
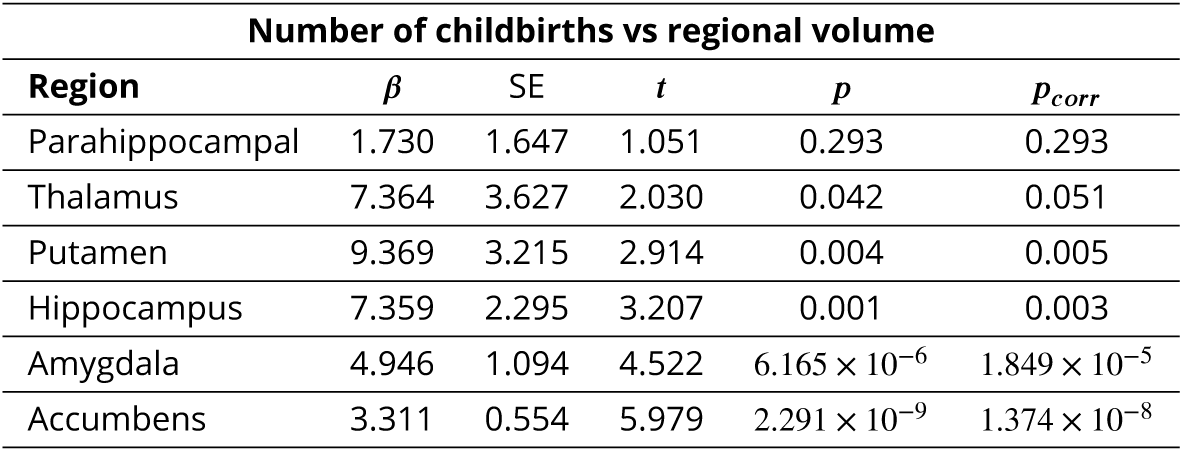
Relationships between number of previous childbirths and volume for each region in subcluster 2. *P*-values are provided before and after FDR correction.

| Number of childbirths vs regional volume |  |  |  |  |  |
| --- | --- | --- | --- | --- | --- |
| Region | $\beta$ | SE | $t$ | $p$ | $p_{corr}$ |
| Parahippocampal | 1.730 | 1.647 | 1.051 | 0.293 | 0.293 |
| Thalamus | 7.364 | 3.627 | 2.030 | 0.042 | 0.051 |
| Putamen | 9.369 | 3.215 | 2.914 | 0.004 | 0.005 |
| Hippocampus | 7.359 | 2.295 | 3.207 | 0.001 | 0.003 |
| Amygdala | 4.946 | 1.094 | 4.522 | $6.165 \times 10^{-6}$ | $1.849 \times 10^{-5}$ |
| Accumbens | 3.311 | 0.554 | 5.979 | $2.291 \times 10^{-9}$ | $1.374 \times 10^{-8}$ |

## Discussion

The results showed that a higher number of previous childbirths was associated with less apparent brain aging in striatal and limbic regions, including the accumbens, putamen, thalamus, hippocampus, and amygdala. The most prominent effect was seen in the accumbens, which is part of the ventral striatum and a key region in the mesolimbic system involved in reward processing and reinforcement learning (***Haber and Knutson, 2010***). The mesolimbic system plays a pivotal role in the rapid emergence of adequate maternal behavior directly after birth due to its role in motivation, reward, and the hedonic value of stimuli (***Brunton and Russell, 2008***; ***Numan and Woodside, 2010***). In rodents, this circuit is activated by pup-related cues that strongly motivate and reinforce maternal care, such as odor (***Fleming et al., 1989***), ultrasonic vocalization (***Robinson et al., 2011***), and suckling (***Ferris et al., 2005***). Low levels of maternal care have been associated with reduced dopamine release within the nucleus accumbens in response to pup cues (***Champagne et al., 2004***), and in humans, motherhood has been linked to anatomical changes in the ventral striatum, with volume reductions promoting responsivity to offspring cues (***Hoekzema et al., 2020***). Together with the ventral striatum, regions including the thalamus, parietal cortex, and brainstem also serve important functions for processing pup-related somatosensory information (***Kim et al., 2010***), and some evidence suggests that structural reorganization occurs in the thalamus, parietal lobe, and somatosensory cortex as a result of physical interactions with pups during nursing (***Kinsley et al., 2008***; ***Xerri et al., 1994***). A recent study by ***Luders et al***. (***2020***) found an increase in regional volumes including the thalamus in women postpartum, corroborating functional MRI (fMRI) studies showing maternal thalamic activation in response to their offspring (***Paul et al., 2019***; ***Rocchetti et al., 2014***). During mother–infant interaction, brain activation has also been shown to increase in the striatum (including putamen and accumbens), amygdala, substantia nigra, insula, inferior frontal gyrus, and temporal gyrus (***Rocchetti et al., 2014***). To summarize, the brain regions identified in the current study largely overlap with neural circuitry underpinning maternal behavior, indicating that brain modulations during pregnancy and postpartum may be traceable decades after childbirth.

In addition to the regions overlapping with the maternal circuit, we found a link between hippocampal brain aging and previous childbirths. This association concurs with animal studies showing enhanced hippocampal neurogenesis in middle age in parous relative to nulliparous rats (***Eid et al., 2019***), and fewer hippocampal deposits of amyloid precursor protein in multiparous relative to primiparous and virgin animals (***Love et al., 2005***). Contrary to the findings in middle-aged animals, *reduced* hippocampal neurogenesis has been reported during the postpartum period, coinciding with enhanced memory performance in primiparous compared to nulliparous rodents (***Kinsley and Lambert, 2008***). In combination with the evidence of both increased and decreased regional volume in humans postpartum (***Luders et al., 2020***; ***Hoekzema et al., 2017***; ***Kim et al., 2018***), these findings emphasize that pregnancy-related brain changes may be highly dynamic.

Pregnancy represents a period of enhanced neuroplasticity of which several underlying mechanisms could confer long-lasting effects on the brain. Fluctuations in hormones including estradiol, progesterone, prolactin, oxytocin, and cortisol are known to influence brain plasticity (***Galea et al., 2014***; ***Simerly, 2002***; ***Barha and Galea, 2010***), and levels of estradiol – a potent regulator of neuroplasticity (***Barha and Galea, 2010***) – rise up to 300-fold during pregnancy (***Schock et al., 2016***) and fall 100–1000 fold postnatally (***Nott et al., 1976***). Hormonal modulations are closely linked to pregnancy-related immune adaptations such as the proliferation of Treg cells (***Kieffer et al., 2017***), which promotes an anti-inflammatory immune environment and contribute to the observed improvement in symptoms of autoimmune disease during pregnancy (***Whitacre et al., 1999***; ***Natri et al., 2019***). In contrast, the transition to menopause marks a period of decline in ovarian hormone levels and can foster a pro-inflammatory phenotype involving increased risk for autoimmune activity and neuronal injury. Beneficial immune adaptations in pregnancy could potentially have long-lasting effects, improving the response to menopause-related inflammation, and subsequently leading to more favorable brain aging trajectories in multiparous women (***Mishra and Brinton, 2018***; ***Fox et al., 2018***; ***Barth and de Lange, 2020***). Another mechanism through which pregnancy may have long-lasting effects on maternal physiology is fetal microchimerism – the presence of fetal cells in the maternal body (***Boddy et al., 2015***). In mice, fetal cells have been found in several brain regions including the hippocampus, where they mature into neurons and integrate into the existing circuitry (***Zeng et al., 2010***). Further evidence for beneficial effects of childbirths on the aging brain stems from studies showing that telomeres are significantly elongated in parous relative to nulliparous women (***Barha et al., 2016***), indicating that parity may slow the pace of cellular aging. However, parity has also been linked to Alzheimer’s-like brain pathology including neurofibrillary tangle and neuritic plaque (***Beeri et al., 2009***; ***Chan et al., 2012***), as well as increased risk of Alzheimer’s disease (***Beeri et al., 2009***; ***Colucci et al., 2006***), with a higher risk in grand-parous women (> 5 childbirths) (***Jang et al., 2018***). Although our previous study showed some evidence of a moderate non-linear trend between number of childbirths and global brain aging (***de Lange et al., 2019***), this effect was not replicated in the current study. More research is needed to determine whether positive effects of pregnancies are less pronounced in grand-parous women, as findings could be biased by low power due to the relatively few women with five or more childbirths, as well as confounding factors such as socioeconomic status or stress levels (***Zeng et al., 2016***).

In conclusion, the current study replicates preceding findings showing less apparent brain aging in multiparous women (***de Lange et al., 2019***), and highlights brain regions that may be particularly influenced by previous childbirths. While prospective longitudinal studies are needed to fully understand any enduring effects of pregnancy, our novel use of regional brain-age prediction – which captures deviations from normative aging – demonstrates that parity relates to region-specific brain aging patterns evident decades after a woman’s last childbirth.

## Methods and Materials

### Sample characteristics

The sample was drawn from the UK Biobank (www.ukbiobank.ac.uk), and included an initial sample of 21,928 women. 1885 participants with known brain disorders were excluded based on ICD10 diagnose (chapter V and VI, field F; *mental and behavioral disorders* and G; *diseases of the nervous system*, except G5 (http://biobank.ndph.ox.ac.uk/showcase/field.cgi?id=41270). 220 participants were excluded based on MRI outliers (see *MRI data acquisition and processing*) and 9 had missing data on number of previous childbirths, yielding a total of 19,787 participants that were included in the analyses. Sample demographics are provided in Table 11.

**Table 11.** Sample demographics. GCSE = General Certificate of Secondary Education, NVQ = National Vocational Qualification.

| **Age** |  |
| --- | --- |
| Mean ± SD | 63.59 ± 7.38 |
| Range [years] | 45.13 - 82.27 |
| **Number of childbirths (live)** |  |
| Mean ± SD | 1.72 ± 1.16 |
| Range | 0 - 9 |
| N in each group: |  |
| **0** = 4297 \| **1** = 2459 \| **2** = 8770 |  |
| **3** = 3334 \| **4** = 729 \| **5** = 142 |  |
| **6** = 43 \| **7** = 7 \| **8** = 5 \| **9** = 1 |  |
| **Age at first birth (N = 15,446)** |  |
| Mean ± SD | 27.08 ± 5.01 |
| Range | 14 - 47 |
| **Years since last birth (N = 13,023)** |  |
| Mean ± SD | 33.37 ± 9.32 |
| Range | 6.77 - 60.34 |
| **Menopausal status (N = 19,781)** |  |
| Yes | 6117 |
| No | 10737 |
| Not sure, had hysterectomy | 1912 |
| Not sure, other reason | 1015 |
| **Ethnic background** |  |
| % White | 97.06 |
| % Black | 0.69 |
| % Mixed | 0.54 |
| % Asian | 0.75 |
| % Chinese | 0.37 |
| % Other | 0.55 |
| % Do not know | 0.03 |
| **Education** |  |
| % University/college degree | 44.71 |
| % A levels or equivalent | 14.09 |
| % O levels/GCSE or equivalent | 24.91 |
| % NVQ or equivalent | 3.17 |
| % Professional qualification | 5.65 |
| % None of the above | 5.91 |
| **Assessment location (imaging)** |  |
| Newcastle | 5139 |
| Cheadle | 11906 |
| Reading | 2742 |

### MRI data acquisition and processing

A detailed overview of the UK Biobank data acquisition and protocols is available in papers by ***Alfaro-Almagro et al***. (***2018***) and ***Miller et al. (2016***). Raw T1-weighted MRI data for all participants were processed using a harmonized analysis pipeline, including automated surface-based morphometry and subcortical segmentation. Volumes of cortical and subcortical brain regions were extracted based on the Desikan-Killiany atlas (***Desikan et al., 2006***) and automatic subcortical segmentation in FreeSurfer (***Fischl et al., 2002***), yielding a set of 68 cortical features (34 per hemisphere) and 17 subcortical features (8 per hemisphere + the brain stem). The MRI data were residualized with respect to scanning site, data quality and motion using Euler numbers (***Rosen et al., 2018***) extracted from FreeSurfer, intracranial volume (***Voevodskaya et al., 2014***), and ethnic background using linear models. To remove bad quality data likely due to subject motion participants with Euler numbers of standard deviation (SD) ± 4 were identified and excluded (n = 192). In addition, participants with SD ± 4 on the global MRI measures mean cortical or subcortical gray matter volume were excluded (n = 13 and n = 22, respectively), yielding a total of 19,796 participants with T1-weighted MRI data. Only participants who had data on number of previous childbirths in addition to MRI were included, and the final sample used in all subsequent analyses (unless otherwise stated) counted 19,787 participants.

### Global and regional brain age prediction

Brain age prediction was used to estimate global and regional brain age based on the MRI data. In line with recent brain-age studies (***de Lange et al., 2020a, 2019***; ***Kaufmann et al., 2019***), the *XG-Boost regressor model*, which is based on a decision-tree ensemble algorithm, was used to run the brain age prediction (https://xgboost.readthedocs.io/en/latest/python). XGboost includes advanced regularization to reduce overfitting (***Chen and Guestrin, 2016***), and uses a gradient boosting framework where the final model is based on a collection of individual models (https://github.com/dmlc/xgboost). Randomized search with ten folds and ten iterations was run to optimize parameters, using all imaging features as input. Scanned parameters ranges were set to *maximum depth*: [2, 10, 1], *number of estimators*: [60, 220, 40], and *learning rate*: [0.1, 0.01, 0.05]. The optimized parameters *maximum depth* = 6, *number of estimators* = 140, and *learning rate* = 0.1 were used for all subsequent models.

To replicate our findings described in ***de Lange et al***. (***2019***), we trained a global brain-age prediction model using the part of the current sample that overlapped with the previous study (N = 10,907), and applied it to the newly added participants (N = 8880), before testing the association between number of previous childbirths and global brain age in the group of new participants. Note that the training set of 10,907 participants overlapping with the previous study showed a lower N relative to the sample in ***de Lange et al***. (***2019***) due to correction for the variables listed in *sample characteristics*.

The full sample (19,787) was utilized to investigate regional brain aging. Averages of the right and left hemisphere measures were first calculated for each feature, and hierarcical clustering was performed on the Spearman rank-order correlation using Scikit-learn version 0.22.2 (https://scikit-learn.org/stable/modules/clustering.html#clustering). All features were grouped according to their associated cluster ID, and separate prediction models were run with ten-fold cross validation, providing cluster-specific brain age gap estimations (predicted age – chronological age) for each individual. To investigate model prediction accuracy, R^2^, RMSE, and MAE were calculated for each model, and correlation analyses were run for predicted versus chronological age. The same procedure was followed for subclusters and region-specific brain age predictions.

### Associations between previous childbirths and regional brain aging

To investigate associations between number of previous childbirths and brain aging, separate regression analyses were run using cluster-specific brain age gap estimates as the dependent variable, and *number of childbirths* as independent variable. Chronological age was included as a covariate, adjusting for age-bias in the brain age predictions as well as age-dependence in number of childbirths (***Le et al., 2018***; ***de Lange and Cole, 2020***). *P*-values were corrected for multiple comparisons using false discovery rate (FDR) correction (***Benjamini and Hochberg, 1995***). To directly compare the associations, *Z* tests for correlated samples (***Zimmerman, 2012***) were run using

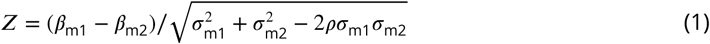

where “m1” and “m2” represent model 1 and 2, the *β* terms represent the beta value from the regression fits, the *σ* terms represent their errors, and *ρ* represents the correlation between the two sets of associations. The statistical analyses were conducted using Python 3.7.0.

## Acknowledgements

This research was conducted using the UK Biobank under Application 27412. While working on this study, the authors received funding from the Research Council of Norway (AM.G.dL.; 286838, T.K.; 276082, L.T.W.; 273345, 249795, 298646, 300768, 223273), the South-East Norway Regional Health Authority (L.T.W.; 2015073, 2019107), the European Research Council under the European Union’s Horizon 2020 research and innovation programme (L.T.W.; 802998), the Alzheimer’s Society (S.S.; Grant Number 441), UK Medical Research Council (K.P.E.; G1001354), and the HDH Wills 1965 Charitable Trust (K.P.E: 1117747).

